# Identification of small molecule inhibitors of G3BP-driven stress granule formation

**DOI:** 10.1101/2023.06.27.546770

**Authors:** Brian D. Freibaum, James Messing, Haruko Nakamura, Ugur Yurtsever, Jinjun Wu, Hong Joo Kim, Jeff Hixon, Rene Lemieux, Jay Duffner, Walter Huynh, Kathy Wong, Michael White, Christia Lee, Rachel Meyers, Roy Parker, J. Paul Taylor

## Abstract

Stress granule formation is triggered by the release of mRNAs from polysomes and is promoted by the action of the paralogs G3BP1 and G3BP2. G3BP1/2 proteins bind mRNAs and thereby promote the condensation of mRNPs into stress granules. Stress granules have been implicated in several disease states, including cancer and neurodegeneration. Consequently, compounds that limit stress granule formation or promote their dissolution have potential as both experimental tools and novel therapeutics. Herein, we describe two small molecules, referred to as G3BP inhibitor a and b (G3Ia and G3Ib), designed to bind to a specific pocket in G3BP1/2 that is known to be targeted by viral inhibitors of G3BP1/2 function. In addition to disrupting co-condensation of RNA, G3BP1, and caprin 1 *in vitro*, these compounds inhibit stress granule formation in cells treated prior to or concurrent with stress, and dissolve pre-existing stress granules when added to cells after stress granule formation. These effects are consistent across multiple cell types and a variety of initiating stressors. Thus, these compounds represent ideal tools to probe the biology of stress granules and hold promise for therapeutic interventions designed to modulate stress granule formation.

## Introduction

Biomolecular condensation underlies the formation of diverse non-membrane-bound intracellular assemblies, including a wide variety of ribonucleoprotein (RNP) granules [1]. Biomolecular condensation can occur through liquid-liquid phase separation (LLPS), a process wherein macromolecular components of a mixed phase de-mix to produce two spatial regions: a higher density, condensed phase containing high concentrations of macromolecules, and a lower density, dilute phase that contains lower concentrations of these macromolecules [2].

RNP granules are one type of biomolecular condensate that can be found throughout the cell, including the nucleus (e.g., nucleoli, Cajal bodies, speckles), cytoplasm (e.g., P bodies, stress granules), and neuronal processes (e.g., RNA transport granules) [3]. Composed of RNA and protein, these condensates vary widely in size and cellular function, with roles that include RNA metabolism, ribosome biogenesis, and signal transduction [1]. On a broader level, the complex and intersecting protein-RNA networks that govern the formation of these condensates are key determinants of intracellular organization [1, 4, 5].

Stress granules are a specialized type of cytoplasmic RNP granule that have been widely used as an archetypal RNP granule to uncover fundamental principles of biomolecular condensation [6, 7]. Stress granules have been implicated in several disease states, including cancer, where they may promote tumor chemoresistance [8–10], and neurodegeneration, where aberrant granule condensation may lead to pleotropic cellular defects and accrual of proteinaceous deposits [7, 11–14]. Thus, targeting stress granules holds promise for therapeutic purposes [15].

Stress granules undergo an ordered assembly and disassembly process driven by a combination of protein-protein, protein-RNA, and RNA-RNA interactions [16, 17] within a defined RNA-protein network [5]. The most central node within this network is the RNA-binding protein G3BP1 and its paralog G3BP2 [5, 18–21]. Depletion of G3BP1/2 from cells eliminates stress granule assembly in response to some stresses [5, 22], and disruption of G3BP1/2 can result in decondensation [19]. For example, ubiquitination of G3BP1 in the context of heat stress enables the selective extraction of G3BP1 from the stress granule network, causing the system to fall below the percolation threshold and disassemble [19]. The reverse is also true: enforced assembly of G3BP1 multimers triggers the formation of stress granules even in the absence of an initiating stress [23], and supplementation of cellular lysates with purified G3BP1 induces concentration-dependent condensation to form granules with the characteristic proteomic and transcriptomic composition of stress granules [24].

G3BP1/2 exists as a dimer, with each monomer comprising a folded dimerization domain (NTF2L), a folded RNA recognition motif (RRM), and three intrinsically disordered regions (IDRs). Dimerization via the NTF2L domain is necessary for stress granule assembly, and experimental disruption of G3BP dimerization results in rapid disassembly of fully formed stress granules [5]. The NTF2L domain also harbors the binding site for other stress granule proteins, including the RNA-binding proteins caprin 1 and USP10, which respectively promote or limit stress granule assembly [22]. Indeed, binding of stress granule proteins at this site may serve to coordinate the interaction network that governs the ordered assembly of stress granules [5].

Insight into the specific properties of NTF2L binding have emerged from unrelated studies examining the role of G3BP1 as a regulator of viral replication [25–27]. For example, Chikungunya viruses inhibit stress granule formation through interaction of the viral nsP3 peptide with a specific binding pocket within the NTF2L domain of G3BP1/2 [28]. Deeper examination of the nsP3 peptide has identified the core G3BP1/2-binding motif as consisting of two FGDF motifs, in which both phenylalanine and the glycine residue are required for binding [26]. This viral strategy may be exploited to modulate human disease, as transfection of nsP3 into mammalian cells can disrupt stress granule formation [29].

Here we designed stable molecules to mimic the interaction of the FGDF motif of nsP3 with G3BP1/2 [30]. We succeeded in developing two molecules, which we refer to as G3BP inhibitor a (G3Ia) and G3BP inhibitor b (G3Ib), that blocked the association of G3BP with its binding partners and inhibited condensation of G3BP both in vitro and in live cells. Treatment of cells with these molecules resulted in the prevention and/or dissolution of stress granules induced either through exogenous stresses or through the expression of disease-causing mutant proteins. These studies demonstrate that the nsP3 binding pocket of G3BP is an effective target to disrupt stress granules in multiple contexts, including a variety of cell types and initiating stressors. Furthermore, the efficacy of our small molecule compounds suggest potential for this approach both in the context of therapeutics and as tools to enable mechanistic studies of RNP granule assembly.

## RESULTS

### Identification of peptide mimetics that bind G3BP1

Previous investigations into the role of G3BP1 as a regulator of viral infections have demonstrated that stress granule formation can be blocked by the binding of the viral nsP3 peptide to the NTF2L domain of G3BP1 [26, 28, 29]. Guided by these studies, we sought to design a series of related modified peptides that could function as small molecule inhibitors of stress granule formation. Beginning with the nsP3 25-mer peptide, we first narrowed the peptide down to an 8-mer that retained the ability to bind the NTF2L domain of G3BP1. Using this minimal 8-mer region, we synthesized multiple lead compounds, which we screened for their capacity to tightly bind the nsP3 binding site within the NFT2L domain of G3BP1 (**Figure S1**). Two such compounds, referred to herein as G3BP inhibitor a and b (G3Ia and G3Ib), bound to G3BP1 with respective K_d_ values of 0.54 μM and 0.15 μM binding as assayed by SPR (**Figure 1A, B**). As negative controls, we also synthesized inactive enantiomers (G3Ia′ and G3Ib′), which bound the same domain greater than 100 folds (K_d_ values of 75.5 μM and 44.5 μM, respectively). To confirm binding to the FGDF pocket of G3BP, we performed a peptide displacement assay such that FRET activity was lost when a compound displaced a FAM-labeled FGDF peptide from the NTF2L domain of G3BP1. We found that G3Ia and G3Ib potently displaced the FGDF peptide, whereas their corresponding enantiomers had little effect, even at very high doses (**Figure 1C**), consistent with the strongly reduced affinity of the enantiomers for the NTF2L domain of G3BP1 (**Figure 1B).** To better understand the nature of the tight binding between G3Ia and the NTF2L domain, we obtained an x-ray crystal structure that revealed a number of important hydrogen bonds between the compound and N122, K123, and F124 of the protein. In addition, a key phenyl ring in G3Ia extends the pi stacking network formed by F15 and F33 of the NTF2L domain (**Figure 1D**). Given the promising properties of G3Ia and G3Ib, including their relatively small polar surface areas suggesting that they might be cell-penetrant (**Figure 1E**), we prioritized G3Ia and G3Ib for further exploration of their capacity to inhibit G3BP-dependent phase separation and/or stress granule formation in cells.

**Figure 1.**
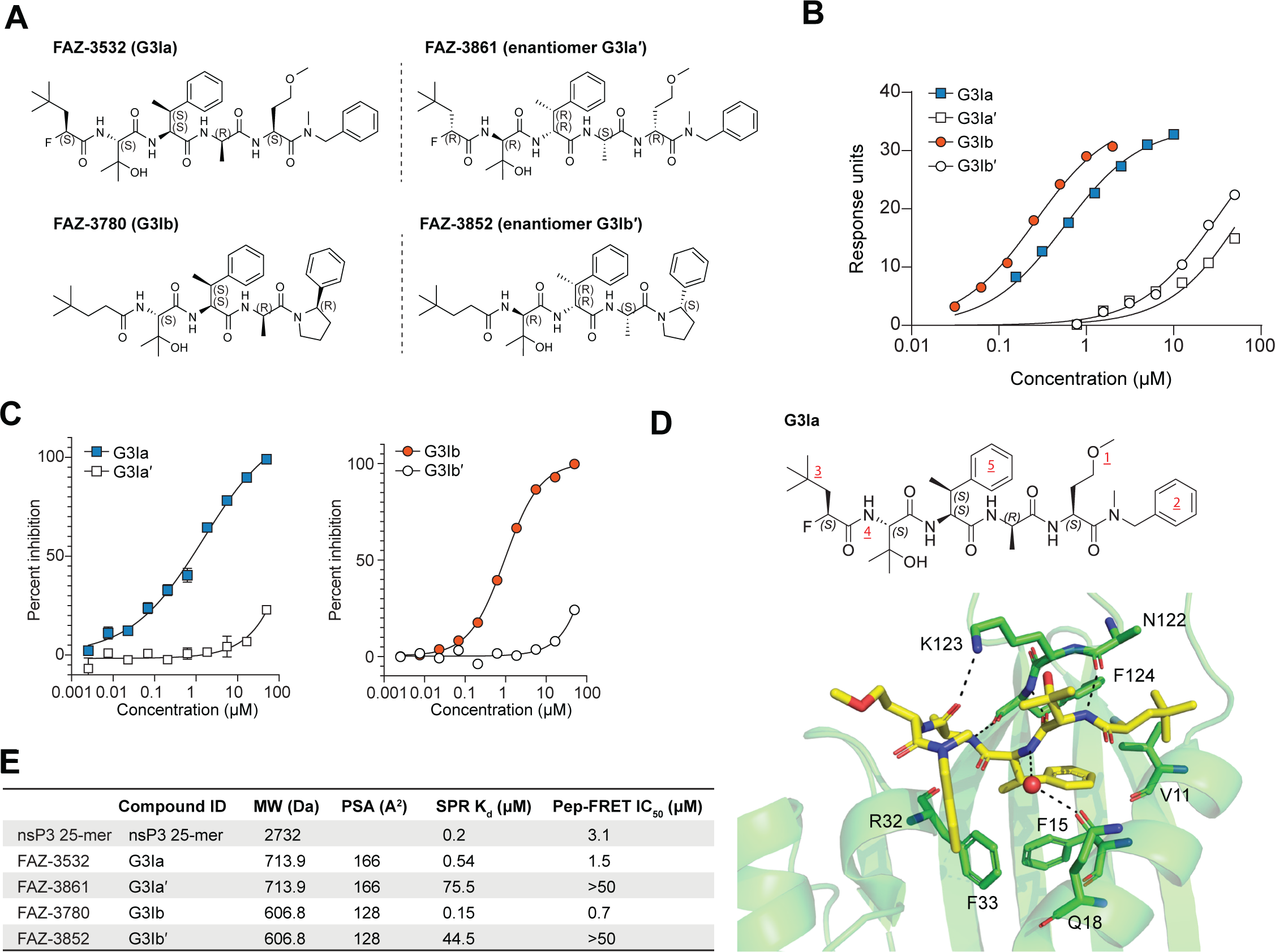
Lead compounds G3Ia and G3Ib bind with high affinity to the NTF2L nsP3 binding pocket of G3BP1. **(A)** Lead compounds FAZ-3532 (G3Ia) and FAZ-3780 (G3Ib) along with respective enantiomer controls FAZ-3861 (G3Ia′) and FAZ-3852 (G3Ib′). **(B)** Representative double-reference subtracted sensorgrams of compounds binding to sensor-immobilized human G3BP1. Compounds were tested by 1/2 dilution with top concentration of 50 μM (G3Ia′), 50 μM (G3Ib′), 10 μM (G3Ia), and 2 μM (G3Ib). Marks on each curve indicate the time span at which equilibrium binding was measured in order to estimate the equilibrium dissociation constant using a 1:1 Langmuir binding model. **(C)** Percent inhibition calculated using the peptide displacement assay at indicated doses of G3I compounds. Error bars represent mean ± SD, n = 2 replicates per dose. **(D)** Crystal structure showing the interaction of G3Ia with the nsP3 binding pocket in the NTF2L domain of G3BP1. The NTF2L domain of G3BP1 (light green cartoon model) crystallized in the presence of G3Ia (yellow sticks), with six copies in the asymmetric unit and copy A shown above. All copies were compound bound, although only half had full compound density. The other three were incomplete in either the ether group (***<u>1</u>***) or terminal phenylalanine (***<u>2</u>***) highlighting their flexibility. Tert-butyl (***<u>3</u>***) functions as a space-filling moiety, maximizing the hydrophobicity of the subpocket lined by V11 and F124. An indirect water-mediated backbone interaction with Q18 is present in four of six copies, including copy A above (large red ball). **(E)** Summary characteristics of the four G3I compounds compared to the nsP3 25-mer peptide. PSA, polar surface area.

### G3Ia and G3Ib disrupt in vitro condensation of RNA, G3BP1, and caprin 1

Binding of the nsP3 peptide to G3BP1 reduces interaction between G3BP1 and its protein partners caprin 1 and USP10 [22]. To test whether G3I compounds had the same effect as nsP3 peptide, we added these compounds to lysates from U2OS cells expressing G3BP1-GFP and used co-immunoprecipitation to assess the interaction between G3BP1 and endogenous caprin1 and USP10. As predicted, addition of G3Ib to these lysates resulted in a dose-dependent reduction of the interaction between G3BP1 and both endogenous caprin 1 and endogenous USP10, whereas the inactive enantiomer G3Ib′ had no such effect (**Figure S2 A-D**). This dose-dependent reduction in interaction with caprin 1 and USP10 was also observed in lysates expressing the NTF2L domain of G3BP1 (GFP-NTF2L) (**Figure S2E, F**).

Caprin 1 is thought to facilitate stress granule assembly by providing additional valency within the multimeric protein-RNA interaction network, thus promoting LLPS and stress granule formation in cells [4, 5]. Indeed, caprin 1, G3BP1, and RNA co-condensate readily in vitro [31], and adding purified recombinant caprin 1 to a mixture of G3BP1 and RNA significantly reduces the threshold concentration of G3BP1 and RNA necessary for LLPS to occur [5]. To test the effect of our compounds on condensate formation by these molecules in vitro, we added G3I compounds to mixtures of purified recombinant caprin 1, G3BP1, and RNA, and found that both G3Ia and G3Ib, but not G3Ia′ or G3Ib′, disrupted the co-condensation of these molecules (**Figure 2A**). These results suggest that active G3I compounds effectively disrupt binding of G3BP1 with caprin 1 and reduce condensation of G3BP1 with RNA. With this mechanism of action in mind, we hypothesized that condensate formation would be unaffected in a 2-component system that consisted of G3BP1 protein and RNA but lacked a binding partner for the NTF2L domain. Indeed, we found that condensate formation by G3BP1 and RNA was unaffected by the addition of G3Ib, even at very high concentrations (166 μM).(**Figure 2B**). In light of these experimental results, the predicted cell permeability of the G3I compounds, and the lack of measurable cell-based toxicity (**Figure 2C**), we pursued these compounds as potential modulators of stress granule formation in living cells.

**Figure 2.**
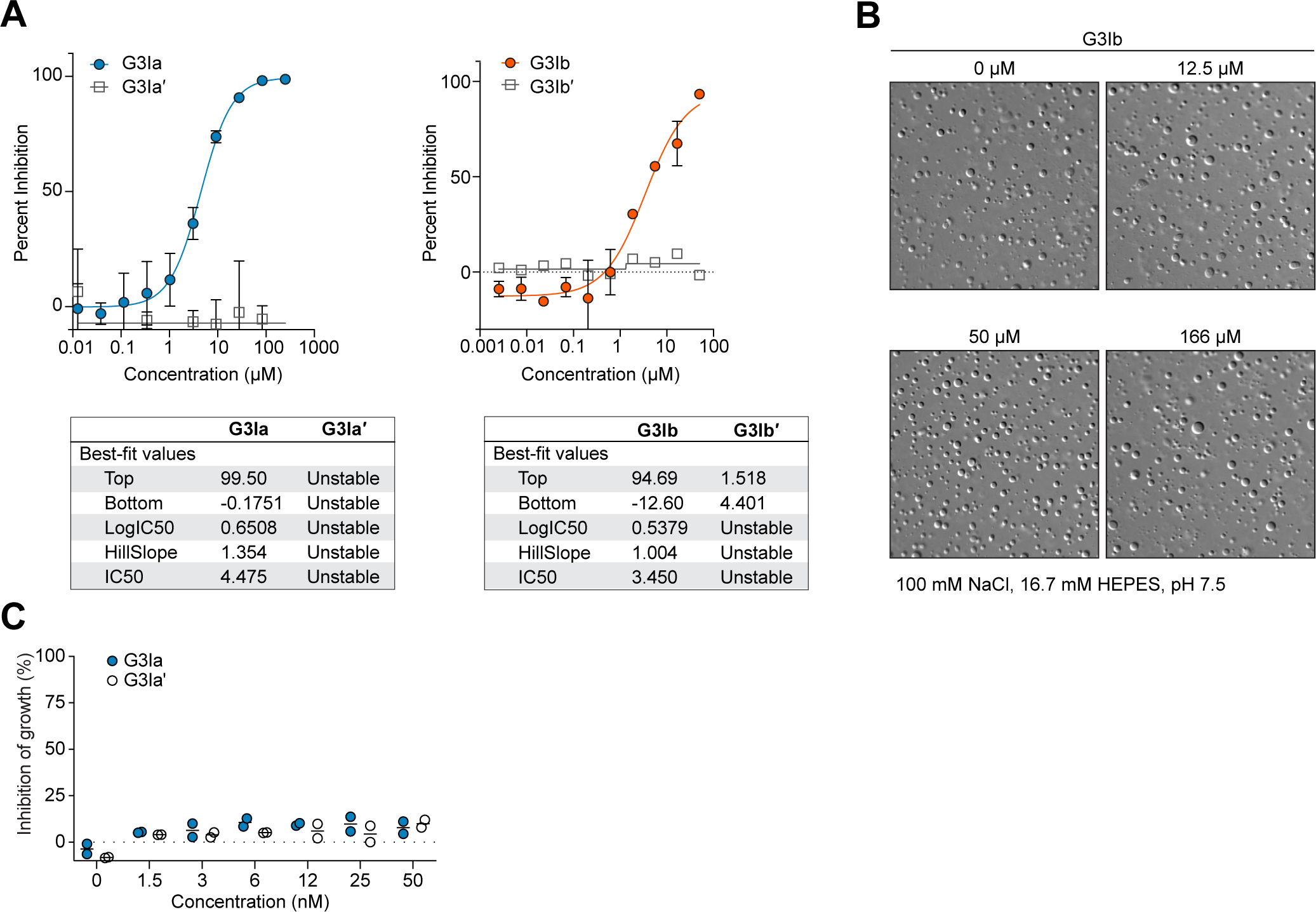
G3Ia and G3Ib disrupt in vitro condensation of RNA, G3BP1, and caprin 1. **(A)** 1.5 μM G3BP1, 1.5 μM caprin 1, and 20 ng/μL total RNA were co-incubated in a three-component system and co-condensation was assessed in the presence of increasing concentrations of G3I compounds. The percent inhibition of G3BP1-GFP in vitro phase separation is shown. Error bars represent mean ± SD. **(B)** 20 μM purified G3BP1 and 100 ng genomic RNA were co-incubated in a two-component system and condensation was assessed in the presence of indicated doses of G3Ib or vehicle control. Condensate formation by G3BP1 and RNA was unaffected by the addition of G3Ib. **(C)** U2OS cells were treated with indicated concentrations of compounds for 24 h and inhibition of growth was measured by monitoring ATP levels, read out through a luciferase signal. N=2, both replicates are plotted.

### Pre-incubation with G3Ia or G3Ib prevents the formation of stress granules in cells

To test the effectiveness of G3Ia and G3Ib in blocking stress granule formation in cells, we used U2OS cells stably expressing G3BP1-GFP and examined the formation of stress granules using live cell imaging with and without these compounds [32]. We began by pre-treating cells with varying concentrations of compounds for 20 minutes, followed by the addition of oxidative stress (500 μM NaAsO_2_) to induce stress granule formation (**Figure 3A**). In cells pretreated with vehicle (DMSO), stress granule formation began approximately 6 minutes after the addition of sodium arsenite, followed by a period of growth and fusion that extended until approximately 20 minutes after the addition of stress (**Figure 3B, C)**.

**Figure 3.**
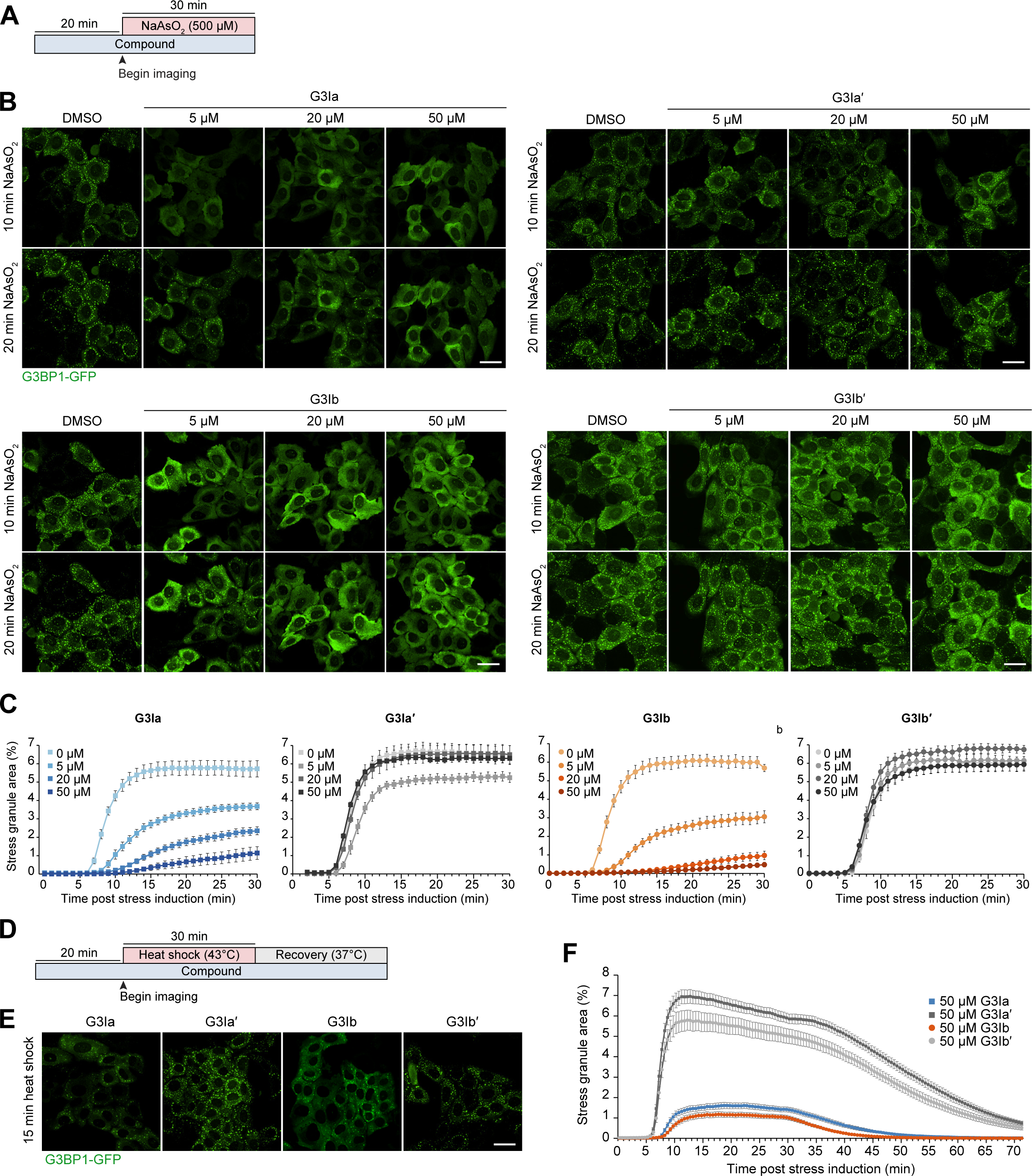
Pre-incubation with G3Ia or G3Ib prevents the formation of stress granules in living cells. **(A)** Schematic showing the pre-incubation paradigm used in panels B-C. Indicated doses of compound were added to cells for 20 min, followed by exposure to 500 μM NaAsO_2_ stress and live cell imaging to monitor stress granule formation. **(B)** Representative images of G3BP1-GFP signal in U2OS cells after 10 min or 20 min of oxidative stress. Scale bars, 40 μm. **(C)** Quantification of cells as in (B) showing the percentage of stress granule area per cell. **(D)** Schematic showing the pre-incubation paradigm used in panels E-F. 50 μM of indicated compounds was added to cells for 20 min, followed by exposure to 43°C heat shock for 30 min. Live cell imaging was used to monitor stress granule formation. **(E)** Representative images of G3BP1-GFP signal in U2OS cells 15 min post heat shock. Scale bar, 40 μm. **(F)** Quantification of cells as in (E) showing the percentage of stress granule area per cell throughout heat shock (43°C, 30 min) and recovery (37°C, 43 min). Error bars represent mean ± SEM in all graphs.

When cells were pretreated with increasing doses of G3Ia or G3Ib, stress granule formation was robustly inhibited in a dose-dependent manner (**Figure 3B, C; Supplementary Videos 1-8**). G3Ia reduced total stress granule area by ∼36% at 5 μM, ∼59% at 20 μM, and ∼80% at 50 μM. G3Ib inhibited stress granule formation even more potently (**Figure 3B, C; Supplementary Videos 1-8**), reducing total stress granule area by ∼50% at 5 μM, ∼85% at 20 μM, and ∼93% at 50 μM. Stress granule formation was not significantly altered by the addition of either of the inactive enantiomers (G3Ia′ or G3Ib′) at any tested dose (**Figure 3B, C; Supplementary Videos 9-16**). Pre-incubation with G3Ia or G3Ib inhibited stress granule formation somewhat more effectively when the concentration of sodium arsenite was reduced by half (250 μM NaAsO_2_), with G3Ia reducing stress granule formation by ∼86% at 50 μM and G3Ib reducing stress granule formation by ∼94% at 50 μM compared to their inactive enantiomers (**Figure S3A-C**).

Because our live-cell assays focused exclusively on G3BP1, it was formally possible that the G3I compounds did not result in inhibition of stress granule assembly, but rather prevented G3BP1 from localizing to otherwise intact stress granules. To test this possibility, we used immunofluorescence to examine additional stress granule markers in cells pretreated with 50 μM G3I compounds and then exposed to 250 μM NaAsO_2_ for 30 minutes. (**Figure S3D**). We found that treatment with G3Ia or G3Ib, but not their inactive enantiomers, resulted in reduced eIF3η and PABPC1 puncta, demonstrating inhibition of stress granule assembly beyond simple extraction or exclusion of G3BP1.

To test whether G3Ia and G3Ib would have comparable effects in other cell lines, we examined their effects on stress granule formation in HeLa cells stably expressing G3BP1-GFP. Similar to our observations in U2OS cells, we found that 50 μM G3Ia and G3Ib inhibited stress granule formation in HeLa cells by ∼72% and ∼82%, respectively, compared with their inactive enantiomers (**Figure S3E, F**).

We also assessed whether the ability of these compounds to block stress granule formation in cells was long lived. To test this, we pretreated U2OS cells with compounds for 24 hours prior to exposure to oxidative stress (500 μM NaAsO_2_ for 30 min). G3Ia exhibited no significant impairment of stress granule formation at concentrations of 5 μM and 20 μM. However, at a concentration of 50 μM, G3Ia inhibited stress granule formation with reduced effectiveness. On the other hand, G3Ib nearly completely inhibited stress granule formation when dosed at 20 μM and 50 μM for 24 hours, indicating that G3Ib has a stronger inhibitory effect on stress granule formation compared to G3Ia under these experimental conditions (**Figure S4**).

The composition and the mechanisms of assembly and disassembly of stress granules can vary based on the type of initiating stress [19, 33, 34]. With this in mind, we next examined whether G3Ia and G3Ib could prevent stress granule formation following heat stress (**Figure 3D**). We found that pre-incubation with G3Ia or G3Ib inhibited stress granule formation, with G3Ia reducing stress granule formation by ∼79% at 50 μM and G3Ib reducing stress granule formation by ∼82% at 50 μM (15 min post heat shock) compared to their inactive enantiomers (**Figure 3E, F**).

Taken together, these results demonstrate that G3Ia and G3Ib effectively prevent stress granule growth and fusion in a dose-dependent manner in multiple cell types and in response to different types of cellular stress.

### Treatment with G3Ia and G3Ib rapidly dissolves pre-formed stress granules

We next tested whether G3Ia and G3Ib could cause the dissolution of pre-formed stress granules. For these experiments, we first treated U2OS cells stably expressing G3BP1-GFP with sodium arsenite (250 μM) for 30 minutes to induce stress granule formation. We then added G3Ia, G3Ib, or their inactive enantiomers, and monitored the cells using live cell imaging. Remarkably, addition of 50 μM G3Ia or G3Ib nearly instantaneously reduced stress granule area by ∼74% and ∼84%, respectively, whereas inactive enantiomers had no statistically significant effect on stress granule area (**Figure 4A, B, Supplementary Videos 17-20**).

**Figure 4.**
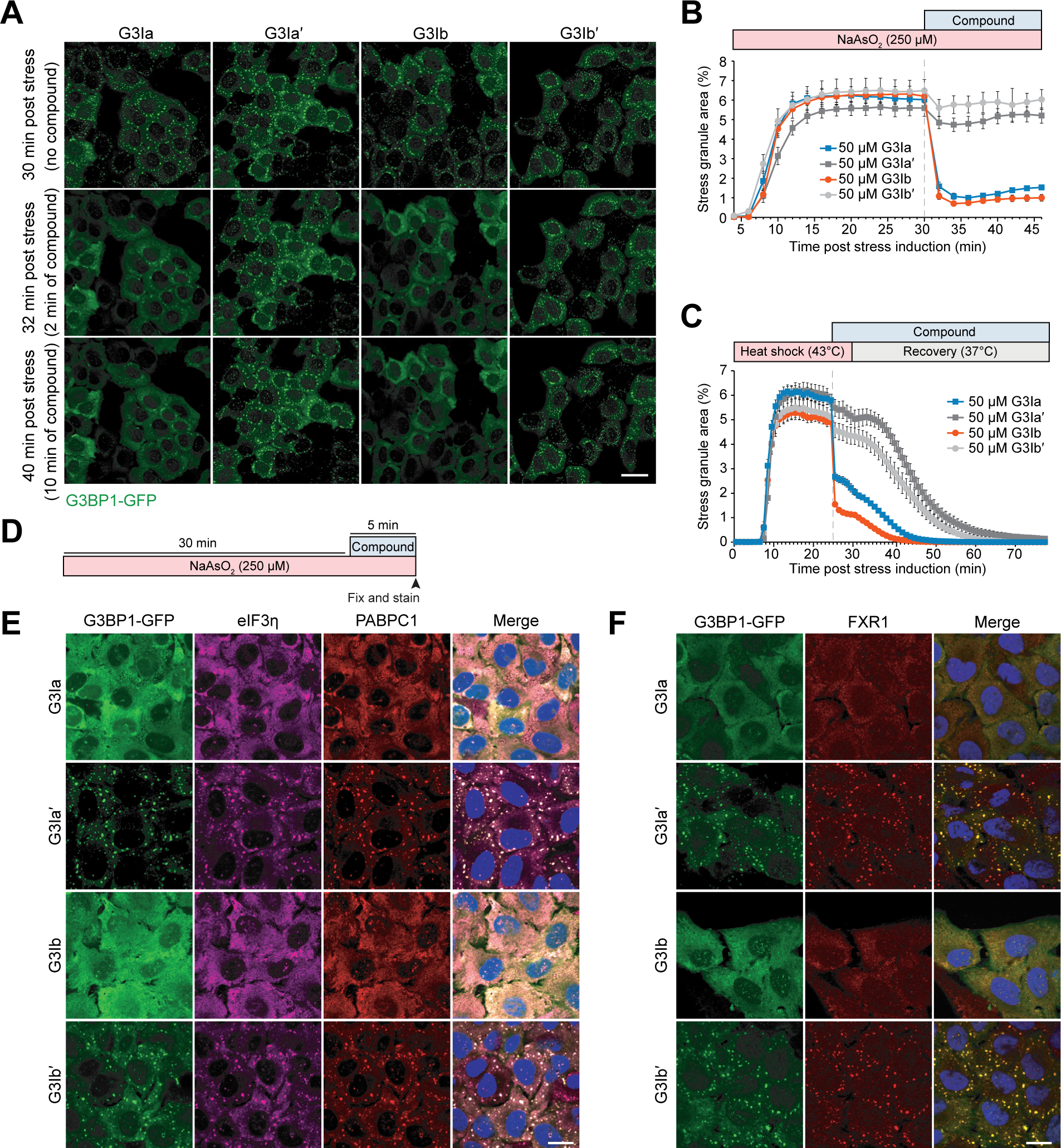
Treatment with G3Ia and G3Ib rapidly dissolves pre-formed stress granules. **(A)** Representative images of G3BP1-GFP signal in U2OS cells following induction of stress by 250 μM NaAsO_2_. Images are shown at 30 min after induction of stress (immediately before addition of compound), 32 min after induction of stress (2 min after addition of 50 μM G3I compound), and 40 min after induction of stress (10 min after addition of 50 μM G3I compound). Scale bar, 40 μm. **(B)** Quantification of cells as in (A) showing the percentage of stress granule area per cell. **(C)** Quantification of the percentage of stress granule area per cell throughout heat shock (43°C, 30 min) and recovery (37°C) from U2OS cells stably expressing G3BP1-GFP. Cells were treated with 50 μM G3I compound 25 min after the induction of heat shock. **(D)** Schematic showing the experimental paradigm used in panels E-F. U2OS cells stably expressing G3BP1-GFP were exposed to 250 μM NaAsO_2_ for 30 min followed by the addition of 50 μM G3I compound. Cells were fixed and stained 5 min after compound was added. **(E-F)** Shown are representative images of immunofluorescence staining of additional stress granule markers (eIF3η, PABPC1, FXR1). Scale bars, 20 μm. Error bars represent mean ± SEM in all graphs.

We also examined whether G3Ia and G3Ib could dissolve stress granules generated through heat stress. Here we exposed cells to 43°C heat stress for 25 minutes, after which we added 50 μM of either G3Ia or G3Ib (**Figure 4C**). Following the addition of G3I compounds, we observed an immediate reduction of ∼54% (G3Ia) and ∼68% (G3Ib) in stress granule area, whereas inactive enantiomers had no immediate effect on stress granule area compared to pre-treatment (**Figure 4C, Supplementary Videos 21-24**). After 5 minutes of incubation with compounds, cells were returned to 37°C, at which point the remaining stress granules began to disassemble in all conditions. As expected, the cells treated with G3Ia or G3Ib completed disassembly faster than cells treated with inactive enantiomers (**Figure 4C, Supplementary Videos 21-24**).

To confirm that the G3I compounds were resulting in disassembly of stress granules, rather than extracting G3BP1 from otherwise intact stress granules, we used immunofluorescence to examine additional stress granule markers in cells exposed to 250 μM NaAsO_2_ for 30 minutes, followed by the addition of 50 μM compound for 5 minutes (**Figure 4D**). We found that treatment with G3Ia or G3Ib, but not their inactive enantiomers, resulted in reduced eIF3η, PABPC1, and FXR1 puncta, indicating disassembly of stress granules beyond simple extraction of G3BP1 (**Figure 4E, F**).

### Treatment with G3Ib prevents the formation of stress granules and dissolves pre-formed stress granules in human iPSC-derived neurons

Given the evidence that aberrant stress granule condensation may contribute to neurodegenerative disease [12], we next tested whether G3I compounds would affect stress granule assembly and/or disassembly in neuronal cells. Here we used iPSC-derived cortical neurons exposed simultaneously to 500 μM NaAsO_2_ and either vehicle, 50 μM G3Ia, or 50 μM G3Ia′ for 60 minutes, at which point we fixed cells and immunostained for G3BP1 (**Figure S5A**). In the absence of stress, ∼3% of neurons had spontaneous stress granules, which increased to ∼70% after 60 minutes of stress (**Figure S5B, C**). Co-incubation with 50 μM G3Ia, but not G3Ia′, blocked both spontaneous and arsenite-induced stress granule formation (**Figure S5B, C**). Next, we examined whether G3I compounds could dissolve pre-formed stress granules in iPSC-derived cortical neurons that had been exposed to 30 minutes of 500 μM NaAsO_2_ stress. Similar to its effects in other cell types, 50 μM G3Ib nearly instantaneously dissolved stress granules in these neuronal cells (**Figure S5D**).

### Treatment with G3Ia or G3Ib dissolves stress granules formed in response to expression of a disease-causing VCP mutant

Stress granules can also form in response to various pathogenic mutations, even in the absence of an exogenous stress. For example, stress granules can arise in cells in response to expression of specific mutant forms of VCP, which in humans lead to the development of multisystem proteinopathy, a condition characterized by degeneration in muscle, bone, and neurons [35]. One such dominant VCP mutation, VCP A232E, induces stress granules in cell culture and also slows the disassembly of stress granules formed through stress [36]. To assess whether G3Ia and G3Ib could alleviate pre-formed granules formed in response to VCP expression, we transfected GFP-VCP A232E into U2OS cells in which endogenous G3BP1 was labeled with tdTomato via CRISPR. Following expression of GFP-VCP A232E, approximately 20% of transfected cells contained stress granules (**Figure 5A**). Adding G3Ia to these cells dissolved 83% of these granules, whereas the inactive enantiomer dissolved only 20% of the granules (**Figure 5A, B**). Similarly, the addition of G3Ib resulted in dissolution of 69% of these granules, in contrast to a reduction of only 6% by G3Ib′ (**Figure 5B**). These observations indicate that G3Ia and G3Ib can dissolve abnormal stress granules that occur due to expression of the pathogenic VCP A232E mutant protein.

**Figure 5.**
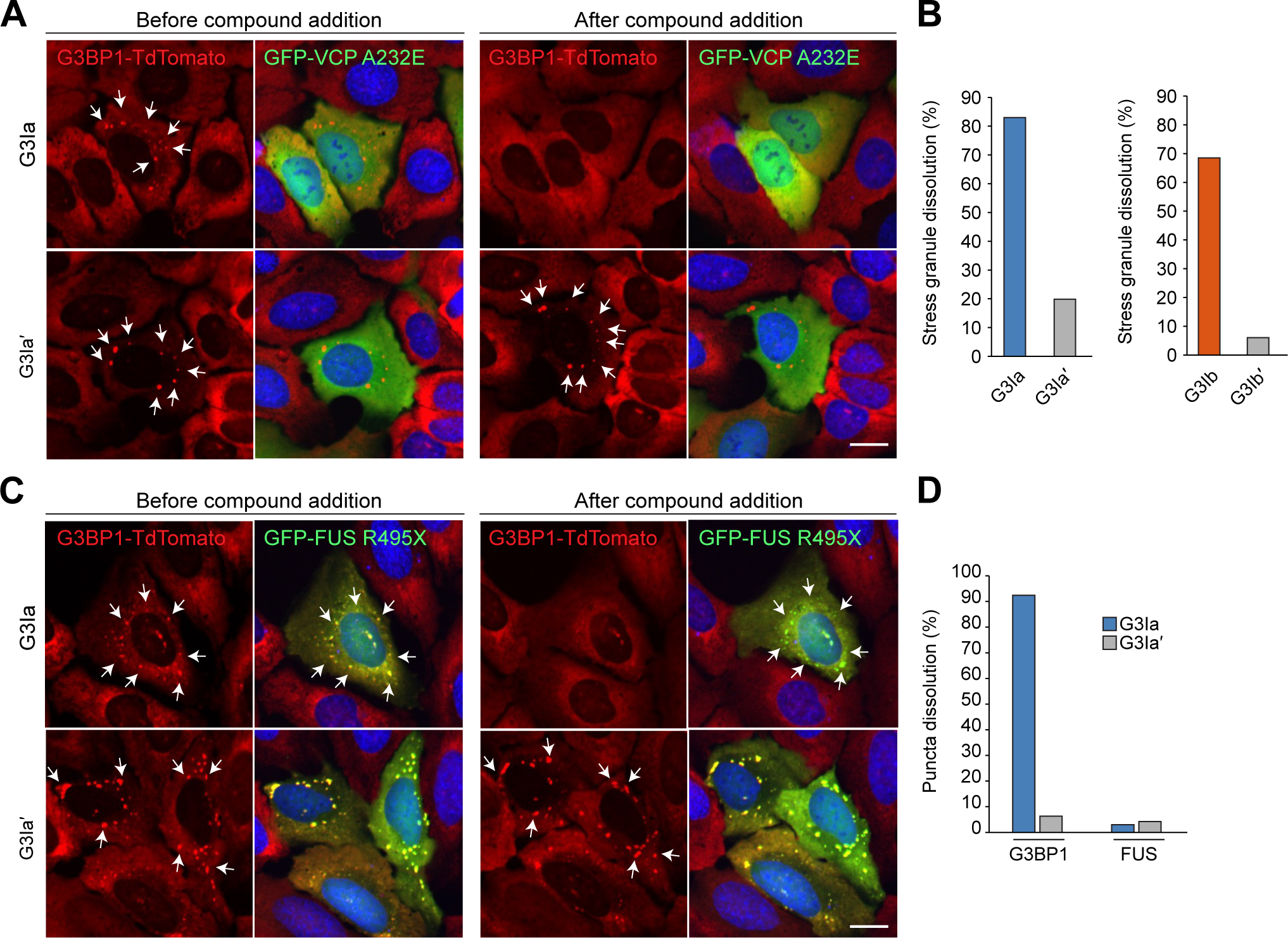
Treatment with G3I compounds modifies stress granules formed in response to expression of disease-causing mutant proteins. **(A)** U2OS cells with TdTomato-tagged endogenous G3BP1 (red) were transfected with VCP A232E (green) for 24 h, then treated with 50 μM G3I compound for 30 minutes. Shown are representative images of cells before (left) and after (right) addition of G3Ia or G3Ia′. Scale bar, 10 μm. **(B)** Quantification of cells as in (A) showing the percentage of stress granule as assessed by TdTomato imaging. Automated puncta tracking was used for G3Ia; manual blinded cell tracking was used for G3Ib. **(C)** U2OS cells with TdTomato-tagged endogenous G3BP1 (red) were transfected with FUS R495X (green) for 24 h, then treated with 50 μM G3I compound for 30 minutes. Shown are representative images of cells before (left) and after (right) addition of G3Ia or G3Ia′. Scale bar, 10 μm. **(D)** Quantification of cells as in (C) showing the percentage of puncta dissolution for G3BP1-positive and FUS R495X-positive puncta.

### Treatment with G3Ia results in removal of G3BP1 from stress granules formed in response to expression of disease-causing FUS mutant

Expression of ALS-causing mutations in the RNA-binding protein FUS leads to the redistribution of mutant FUS protein into the cytoplasm, where it accumulates in stress granules [37]. To test the ability of our compounds to dissolve FUS-containing granules, we transfected the disease-causing mutant FUS R495X into U2OS cells expressing endogenous G3BP1 tagged with tdTomato as described above, and then used automated imaging analysis to examine these cells before and after addition of G3I compounds. Before the addition of compound, approximately 40% of cells expressing FUS R495X contained stress granules in which G3BP1 was co-localized with FUS 495X (**Figure 5C**). Addition of G3Ia eliminated 92% of the G3BP1 puncta, whereas the inactive enantiomer eliminated only 6% of G3BP1 puncta. In contrast, neither G3Ia (3%) nor G3Ia′ (4%) were effective in reducing the number of puncta positive for FUS R495X (**Figure 5C, D**). These results suggest that the FUS inclusions are more stable than typical stress granules and are not dependent on G3BP1 for their persistence.

## DISCUSSION

Using the nsP3 peptide as a lead compound (**Figure 1**), we designed two novel compounds: G3Ia and G3Ib that inhibit the formation of stress granules when added to cells prior to either arsenite or heat stress (**Figure 3**). Additionally, G3Ia and G3Ib trigger the disassembly of existing stress granules when introduced to cells following stress granule formation (**Figure 4**). The presumed mechanism of action of these compounds is through antagonism of binding between G3BP1/2 and their binding partners within the NTF2L domain. Indeed, we have demonstrated that G3Ia and G3Ib inhibit the interaction of G3BP1 with caprin 1 and USP10 (**Figure S1**). The loss of these protein interactions within the NTF2L domain is predicted to limit oligomerization of the G3BP1/2 proteins, thus preventing the valency necessary for the formation of large multimeric protein-RNA complexes [4] that comprise the liquid condensates know as stress granules. Both G3Ia and G3Ib behaved as expected, leading to the inhibition of *in vitro* condensate formation induced through the interaction of G3BP1, caprin 1 and RNA (**Figure 2**).

The use of G3Ia and G3Ib to target the NTF2L domain of G3BP1/2 has several potential applications. Primarily, these compounds will provide a tool for researchers to manipulate G3BP1/2 dependent stress granule formation. Unlike many other stress granule inhibitors such as cycloheximide [38], that lead to the global inhibition of translation and are highly toxic, G3Ia and G3Ib are designed to be highly specific in their inhibition of protein binding within the NTF2L domain of G3BP1/2. G3Ia and G3Ib impair condensate formation in a highly tractable manner, but they do not appear to induce cellular toxicity or growth phenotypes (**Figure 1**) and appear to be highly versatile among different cell types. Additionally, G3Ia and G3Ib only target the protein interaction domain G3BP1/2, thereby allowing the researcher to block G3BP1/2 condensate formation, at the same time leaving intact the dimerization and RNA-binding capabilities of G3BP1/2. This is a significant advantage over the time-consuming genetic models in which G3BP1/2 are knocked out, resulting in a complete loss of function of these proteins in cells. These compounds will likely prove to be highly useful in probing what specific function stress granules perform in cells.

Stress granules have been implicated in several diseases from neurodegenerative diseases to cancer [39]. These compounds could be utilized in disease models to either block stress granule formation entirely or to possibly dissolve disease-initiated granules, allowing for a better understanding of what role these condensates play in either disease progression or prevention. Consistent with this possibility, we observed that G3Ia and G3Ib can induce the disassembly of aberrant stress granules triggered by expression of a pathogenic A232E VCP mutation. This mutation leads to multisystem proteinopathy in patients: resulting in disease of neurons, muscle and bone (**Figure 5**). Additionally, G3Ia and G3Ib can be used to probe the immunological role of stress granules, as G3BP1 mediated stress granules are known to form in response to infection with numerous viral RNAs [40]. Finally, the compounds can used to determine whether the NTF2L domain-mediated interaction of G3BP1/2 plays a role in the normal function of cells outside of its ability to drive condensation. This could lead to discovering novel stress granule independent functions of G3BP1, G3BP2 or their interactors in normal cell biology. In conclusion, G3Ia and G3Ib are potent inhibitors of binding to the NTF2L domain of G3BP1/2 that are likely highly specific, easy to synthesize, and highly tractable, providing a valuable new tool in the study of condensate biology.

---

**Figure S1.**
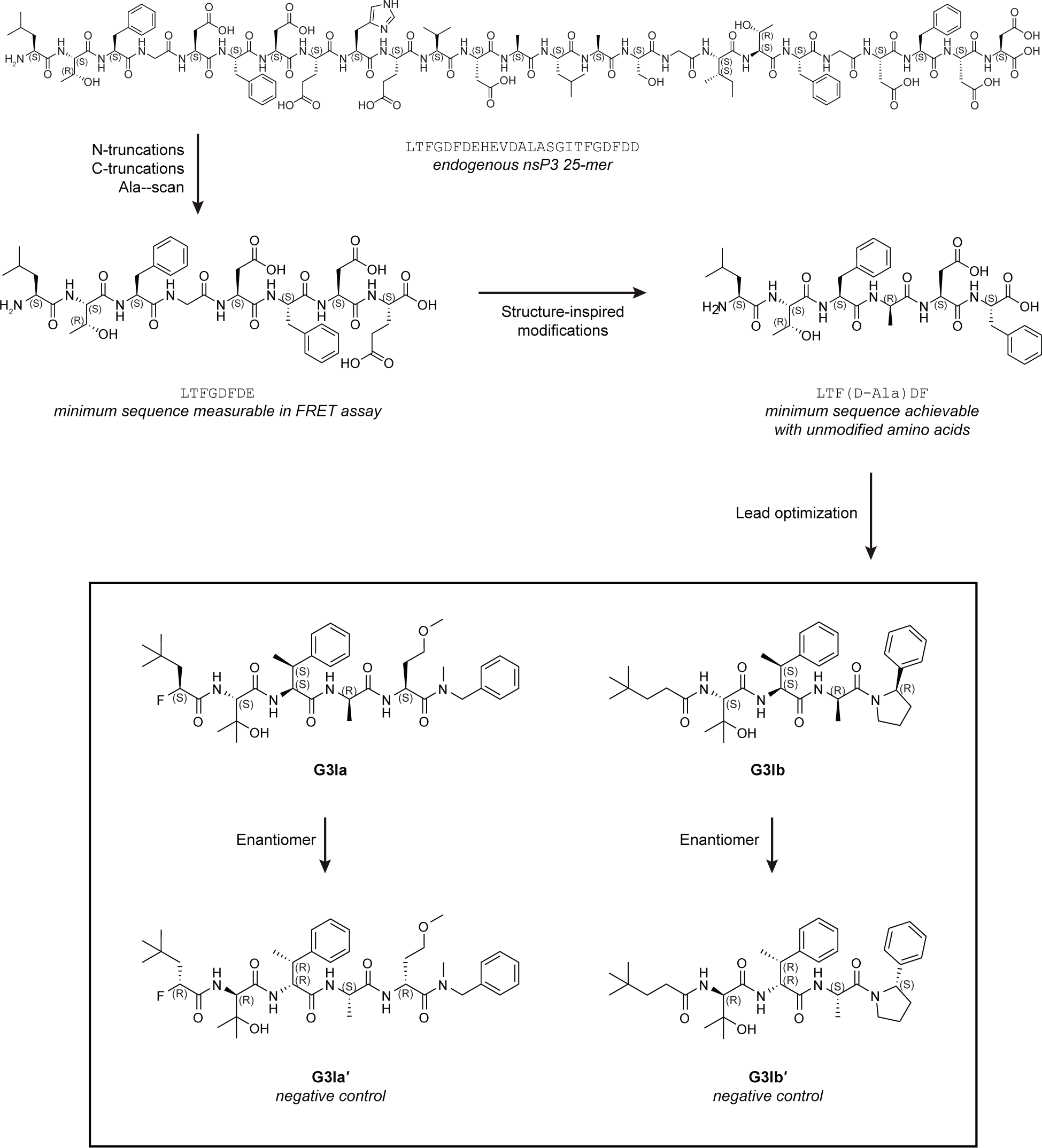
Identification of lead compounds that interact with the nsP3 binding pocket of the NTF2L domain of G3BP1. Schematic showing the lead optimization and compound discovery process that led to the identification of G3Ia and G3Ib as lead compounds.

**Figure S2.**
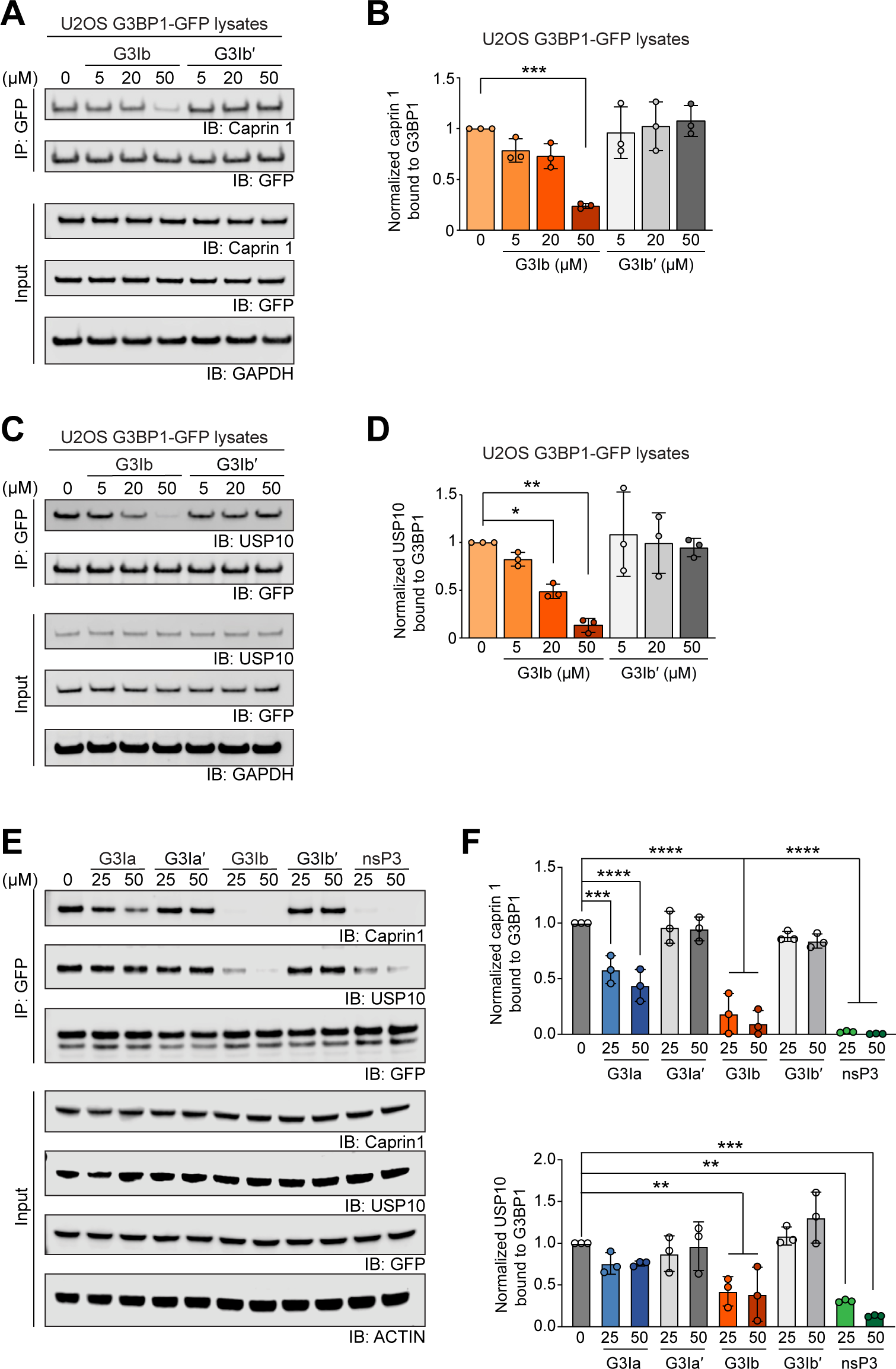
G3Ib disrupts interaction between G3BP1 and its interaction partners caprin 1 and USP10 in a dose-dependent manner. **(A-D)** Lysates from U2OS cells stably expressing G3BP1-GFP were collected, incubated with increasing concentrations of G3I compound, immunoprecipitated for GFP, and separated by SDS-PAGE. Blots were probed for GFP and endogenous caprin 1 (A-B) or endogenous USP10 (C-D). GAPDH was used a loading control. Densitometry from n = 3 blots was used to generate graphs (B) and (D); error bars represent mean ± SD. \**P* = 0.0495, \*\**P* = 0.0011, \*\*\**P* = 0.0002 by one-way ANOVA with Dunnett’s multiple comparisons test. **(E)** Lysates from HEK293T cells expressing GFP-NTF2L were collected, incubated with increasing concentrations of G3I compounds or nsP3, immunoprecipitated for GFP, and separated by SDS-PAGE. Blots were probed for GFP and endogenous caprin 1 or endogenous USP10. Actin was used a loading control. A representative blot is shown from n=3 experiments **(F)** Quantification of densitometry from n = 3 blots. Error bars represent mean ± SD. \*\*\**P* = 0.0006 and \*\*\*\**P* < 0.0001 for caprin 1, \*\**P* = 0.0087 (25 μM G3Ib), \*\**P* = 0.0050 (50 μM G3Ib), \*\**P* = 0.0015 (25 μM nsP3), \*\*\**P* = 0.0001 (50 μM nsP3) for USP10 by one-way ANOVA with Dunnett’s multiple comparisons test.

**Figure S3.**
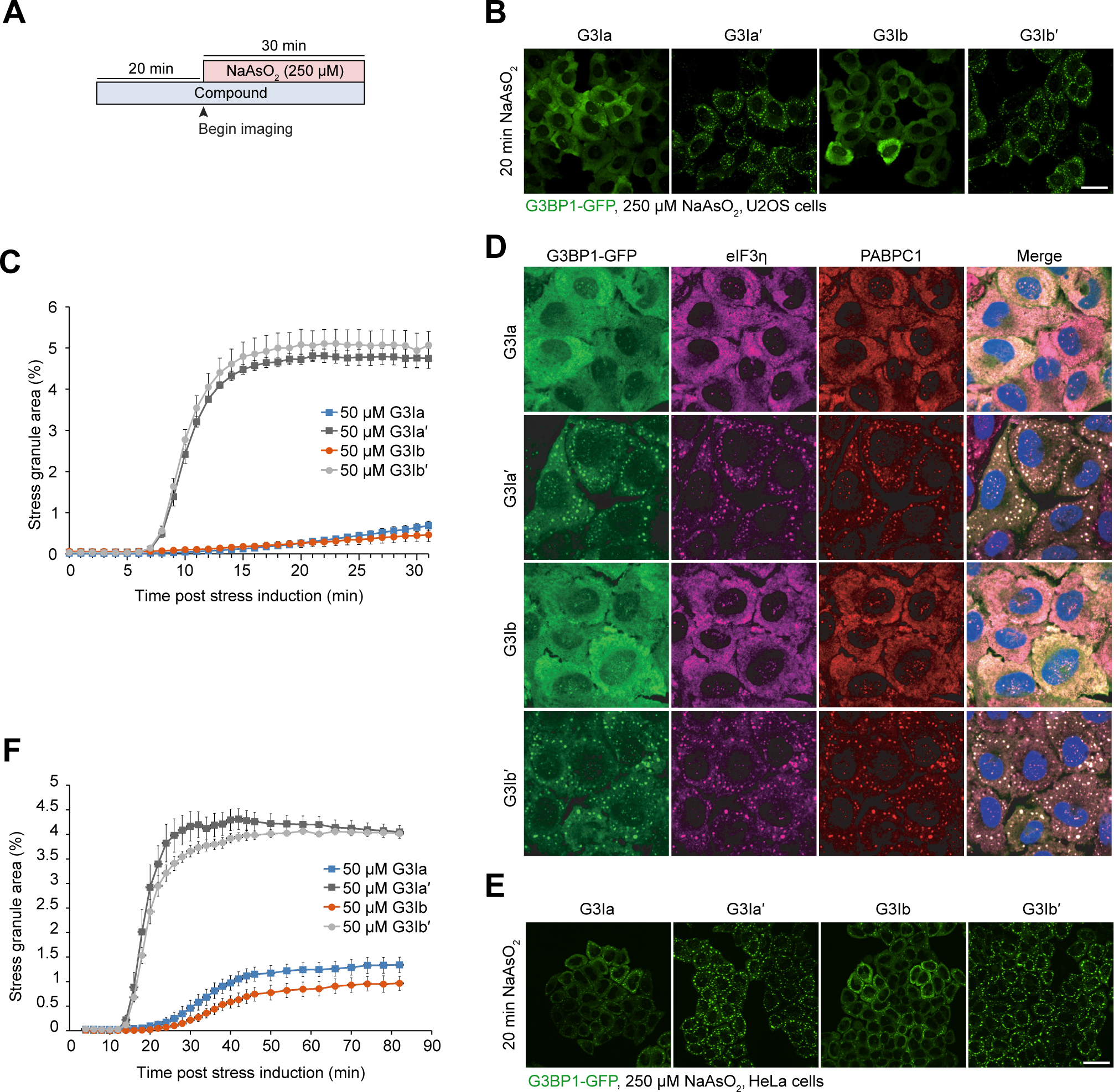
Pre-incubation with G3Ia or G3Ib prevents the formation of stress granules at a lower dose of sodium arsenite and in HeLa cells. **(A)** Schematic showing the pre-incubation paradigm used in panels B-D. 50 μM of indicated compounds was added to cells for 20 min, followed by exposure to 250 μM NaAsO_2_ stress and live cell imaging to monitor stress granule formation. **(B)** Representative images of G3BP1-GFP signal in U2OS cells after 20 min 250 μM NaAsO_2_. Scale bar, 40 μm. **(C)** Quantification of cells as in (B) showing the percentage of stress granule area per cell. **(D)** Representative images of immunofluorescent staining of additional stress granule markers (eIF3η, PABPC1). Scale bar, 20 μm. **(E)** Representative images of G3BP1-GFP signal in HeLa cells after 20 min of 250 μM NaAsO_2_. Scale bar, 40 μm. **(F)** Quantification of cells in (E) showing the percentage of stress granule area per cell. Error bars represent mean ± SEM in all graphs.

**Figure S4.**
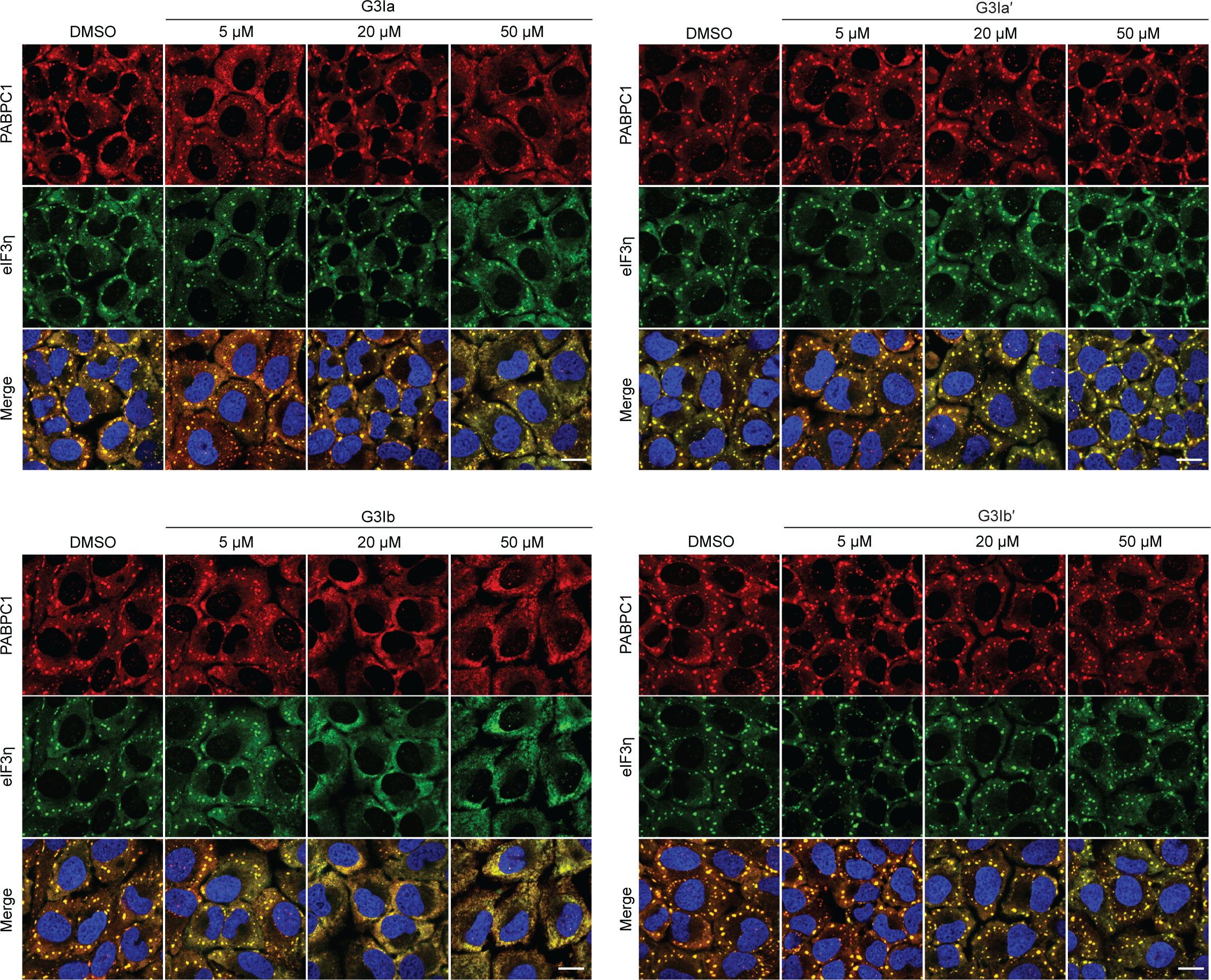
G3Ia or G3Ib continues to inhibit the formation of stress granules 24 hours after exposure to compounds. Indicated doses of compound were added to cells for 24 hours, followed by exposure to 500 μM NaAsO_2_ stress for 30 min to monitor stress granule formation. Shown are representative images of immunofluorescence staining of additional stress granule markers (eIF3η, PABPC1). Scale bars, 40 μm.

**Figure S5.**
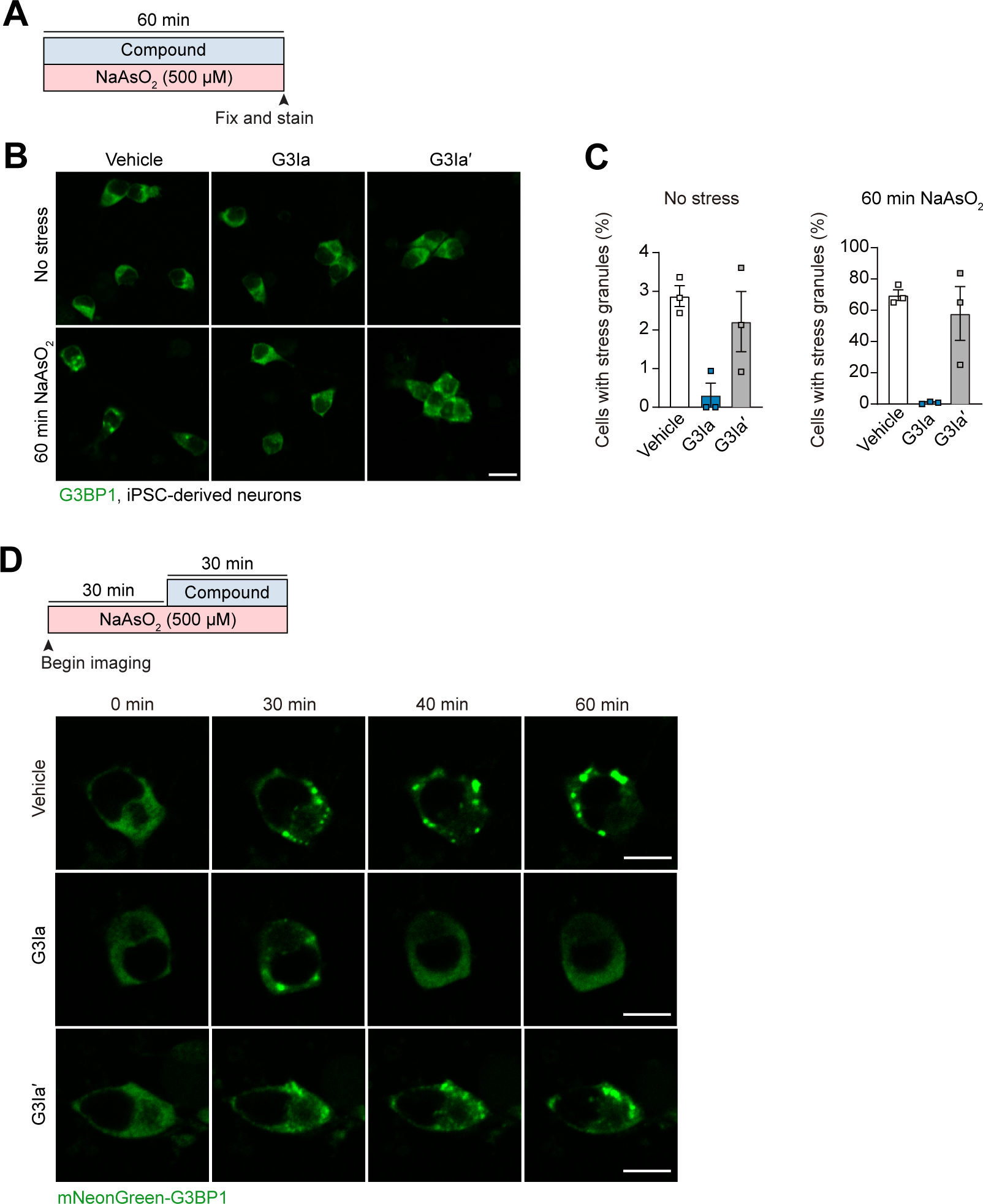
Treatment with G3Ib prevents the formation of stress granules and dissolves pre-formed stress granules in human iPSC-derived neurons. **(A)** Schematic showing the paradigm used in panels B-C. iPSC-derived cortical neurons were treated with 50 μM G3Ia or G3Ia′ in the presence or absence of 500 μM NaAsO_2_ for 60 min followed by fixation and imaging. **(B)** Representative images showing stress granules via staining of G3BP1 in cortical neurons in the presence or absence of G3I compounds under baseline or stressed conditions. **(C)** Quantification of cells as in (B) showing the percentage of cells with stress granules under each condition. Scale bars, 20 μm; error bars represent mean ± SEM. **(D)** Schematic showing G3BP1-mNeonGreen iPSC-derived neurons exposed to 500 μM NaAsO_2_ for 30 min, at which point indicated 50 μM G3I compounds were added. Shown are representative images showing live cell imaging of stress granules via mNeonGreen-tagged G3BP1. Scale bars, 10 μm.

**Supplemental Video 1.** Live cell imaging following pre-incubation with DMSO (control for G3Ia) followed by exposure to 500 μM NaAsO_2_ for 30 min.

**Supplemental Video 2.** Live cell imaging following pre-incubation with 5 μM G3Ia followed by exposure to 500 μM NaAsO_2_ for 30 min.

**Supplemental Video 3.** Live cell imaging following pre-incubation with 20 μM G3Ia followed by exposure to 500 μM NaAsO_2_ for 30 min.

**Supplemental Video 4.** Live cell imaging following pre-incubation with 50 μM G3Ia followed by exposure to 500 μM NaAsO_2_ for 30 min.

**Supplemental Video 5.** Live cell imaging following pre-incubation with DMSO (control for G3Ib) followed by exposure to 500 μM NaAsO_2_ for 30 min.

**Supplemental Video 6.** Live cell imaging following pre-incubation with 5 μM G3Ib followed by exposure to 500 μM NaAsO_2_ for 30 min.

**Supplemental Video 7.** Live cell imaging following pre-incubation with 20 μM G3Ib followed by exposure to 500 μM NaAsO_2_ for 30 min.

**Supplemental Video 8.** Live cell imaging following pre-incubation with 50 μM G3Ib followed by exposure to 500 μM NaAsO_2_ for 30 min.

**Supplemental Video 9.** Live cell imaging following pre-incubation with DMSO (control for G3Ia’) followed by exposure to 500 μM NaAsO_2_ for 30 min.

**Supplemental Video 10.** Live cell imaging following pre-incubation with 5 μM G3Ia’ followed by exposure to 500 μM NaAsO_2_ for 30 min.

**Supplemental Video 11.** Live cell imaging following pre-incubation with 20 μM G3Ia’ followed by exposure to 500 μM NaAsO_2_ for 30 min.

**Supplemental Video 12.** Live cell imaging following pre-incubation with 50 μM G3Ia’ followed by exposure to 500 μM NaAsO_2_ for 30 min.

**Supplemental Video 13.** Live cell imaging following pre-incubation with DMSO (control for G3Ib’) followed by exposure to 500 μM NaAsO_2_ for 30 min.

**Supplemental Video 14.** Live cell imaging following pre-incubation with 5 μM G3Ib’ followed by exposure to 500 μM NaAsO_2_ for 30 min.

**Supplemental Video 15.** Live cell imaging following pre-incubation with 20 μM G3Ib’ followed by exposure to 500 μM NaAsO_2_ for 30 min.

**Supplemental Video 16.** Live cell imaging following pre-incubation with 50 μM G3Ib’ followed by exposure to 500 μM NaAsO_2_ for 30 min.

**Supplemental Video 17.** Live cell imaging following exposure to 250 μM NaAsO_2_ for 30 min followed by treatment with 50 μM G3Ia.

**Supplemental Video 18.** Live cell imaging following exposure to 250 μM NaAsO_2_ for 30 min followed by treatment with 50 μM G3Ia’.

**Supplemental Video 19.** Live cell imaging following exposure to 250 μM NaAsO_2_ for 30 min followed by treatment with 50 μM G3Ib.

**Supplemental Video 20.** Live cell imaging following exposure to 250 μM NaAsO_2_ for 30 min followed by treatment with 50 μM G3Ib’.

**Supplemental Video 21.** Live cell imaging following exposure to a 43°C heat shock for 30 min followed by treatment with 50 μM G3Ia 25 minutes after exposure to heat shock.

**Supplemental Video 22.** Live cell imaging following exposure to a 43°C heat shock for 30 min followed by treatment with 50 μM G3Ia’ 25 minutes after exposure to heat shock.

**Supplemental Video 23.** Live cell imaging following exposure to a 43°C heat shock for 30 min followed by treatment with 50 μM G3Ib 25 minutes after exposure to heat shock.

**Supplemental Video 24.** Live cell imaging following exposure to a 43°C heat shock for 30 min followed by treatment with 50 μM G3Ib’ 25 minutes after exposure to heat shock.

## Methods

### Surface plasmon resonance (SPR)

Binding affinity for G3BP1 was measured by SPR assay using a Biacore 8K instrument (Cytiva) at 25°C at pH 7.4 (20 mM Tris, 300 mM NaCl, 0.05% Tween-20, 2% DMSO) using neutravidin sensor chips. Binding of solution phase molecules was measured to sensor-bound biotinylated, AVI-tagged versions of the NTF2L domain of G3BP1. G3BP1 was captured on the sensor surface at the beginning of each experiment at a level of approximately 1,000 RU. A neutravidin-only surface was used as a reference. Binding affinity was measured by equilibrium analysis of double-reference subtracted data performed at seven different analyte concentrations. A 1:1 binding model was used to estimate affinity; maximum binding response was typically about 90% of the theoretical maximum for a 1:1 interaction given the size of the protein and analytes. Binding at the end of a 90-second association phase was measured; data was only used where equilibrium binding was reached. No regeneration was performed between binding cycles as all tested molecules dissociated completely from the sensor at the end of each cycle. Estimation of binding affinity by kinetic analysis was not performed as kinetic constants were beyond the range that could be reliably measured by the instrumentation.

### Peptide displacement

Peptide displacement assays were performed in 20 μl total volume at 25°C at pH 7.4 in buffer containing 20 mM Tris-HCl, pH 7.5, 100 mM NaCl, 0.01% bovine serum albumin, 1 mM DTT, and 0.005% Tween-20. 10 nM full-length 6x-His-tagged human G3BP1 produced in mammalian cells was added to the reaction mix containing test compound and 0.5 nM anti-His-Tb antibody (PerkinElmer). Assays were initiated by adding 4 nM FAM-labeled PEG6-USP10 24-mer peptide probe (FAM-PEG6-GALHSPQYIFGDFSPDEFNQFFVT; Peptide 2.0). Assay plates were shaken for 30 sec, then centrifuged for 30 sec, then incubated at room temperature for 60 min. Data was read on an EnVision Multilabel plate reader at 495 nm/520 nm. HTRF ratios were calculated as described by PerkinElmer and IC_50_s were calculated using GraphPad Prism 9 software and the [Inhibitor] vs. Response – Variable Slope (four parameter) fit equation.

### Liquid-liquid phase separation

LLPS assays were performed in 30 μl total volume at 25°C at pH 7.4 in buffer containing 37 mM Tris, pH 7.5, 116 mM NaCl, 0.33% NP-40, 2.6% glycerol, 0.5 mM DTT with 20 ng/μL total cellular RNA from U2OS cells and 1.425 μM G3BP1, 0.075 μM FLAG-GFP-G3BP1, and 1.5 μM caprin-1. Reactions were initiated by adding G3BP1/caprin protein mix to cellular RNA, followed by 3 min of shaking at 450 rpm and then incubation for 70 min in the dark. For condensates formed with G3BP1 and RNA, 20 μM purified G3BP1 and 100 ng of genomic RNA was added to a buffer consisting of 100 mM NaCl and 16.7 mM HEPES (pH 7.5) in the presence of the indicated dose of G3Ib or vehicle control.

### Cell culture and transfection

U2OS (HTB-96), HEK293T (CRL-3216) and HeLa (CCL-2) cells were originally purchased from ATCC and periodically authenticated by short tandem repeat (STR) profiling. Cells were grown in Dulbecco’s modified Eagle’s medium (DMEM) supplemented with 10% fetal bovine serum (FBS), 1% penicillin/streptomycin, and 1% L-glutamate. Cells were counted using ADAM-CellT (NanoEntek), plated and transfected using either Lipofectamine 3000 (Thermo Fisher; L3000008) or Viafect (Promega; E4981) for transient overexpression according to the manufacturer’s instructions. Cytotoxicity was assayed using the CellTiter-Glo 2.0 reagent (Promega) according to the manufacturer’s instructions.

### Pre-incubation of compounds and live cell imaging

For pre-incubation of compounds, U2OS or HeLa cells stably expressing G3BP1-GFP were seeded into eight-well Lab-Tek chambered cover glass (Nunc). At least 20 min prior to the experiment, 1 mL FluoroBrite DMEM media (Gibco) supplemented with 10% FBS and 4 mM L-glutamine was combined with 0.1% DMSO or G3I compound (5, 10, or 50 μM). This solution was vortexed for 5 sec, added to the cells at 250 μL per well, and left to incubate on the microscope for 20 min. Conditions were maintained at 37°C and 5% CO_2_ using a Bold Line Cage Incubator (Okolab) and an objective heater (Bioptechs). During this incubation time, 5 xy fields were stored per condition, with each field having ∼20-30 cells within it. After the 20-min incubation, 250 μL 500 μM or 1 mM (2x) sodium arsenite (Sigma) diluted in the FluoroBrite media with an appropriate amount of G3I compound or DMSO was added to the sample. Imaging began immediately after, with images at each xy field being taken every 1 min. For heat shock experiments, U2OS G3BP1-GFP-expressing cells were seeded into a 35-mm glass bottom dish (MatTek). Prior to the experiment, 1.5 mL FluoroBrite DMEM media (Gibco) supplemented with 10% FBS and 4 mM L-glutamine was combined with 0.1% DMSO or 50 μM G3I compound. The sample was incubated on the environmentally controlled microscope for 20 min. During this incubation time, 10 xy fields were stored per condition. After the 20-min incubation, acquisition began with each xy position being imaged every 30 sec. Two min into the acquisition, the objective heater was ramped to 43°C and maintained this temperature until ramped back down to 37°C at 32 min into the acquisition. All imaging was acquired on a Yokogawa CSU W1 spinning disk attached to a Nikon Ti2 eclipse with a Photometrics Prime 95B camera using Nikon Elements software (v5.20.00 to v5.21.02). Imaging was performed through a Nikon Plan Apo 60× 1.40 NA oil objective with Immersol 518 F (Zeiss; refractive index 1.518), and Perfect Focus 2.0 (Nikon) was engaged for all captures.

### Post-addition of compounds and live cell imaging

For post-addition of compounds, U2OS stably expressing G3BP1-GFP were seeded into eight-well Lab-Tek chambered cover glass (Nunc) for sodium arsenite experiments or 35-mm glass-bottom dishes (MatTek) for heat shock experiments. Immediately prior to the experiment, media was replaced with FluoroBrite DMEM media (Gibco) in each well supplemented with 10% FBS and 4 mM L-glutamine; for sodium arsenite experiments, arsenite was added at a final concentration of 250 μM. Cells were then placed on a microscope on which 37°C temperature and 5% CO_2_ were maintained using a Bold Line Cage Incubator (Okolab) and an objective heater (Bioptechs), and 5-10xy fields were stored. For sodium arsenite experiments, acquisition began immediately after finding all xy positions, 4 min after addition of sodium arsenite. Thirty min after arsenite was added, 250 μL FluoroBrite DMEM with 250 μM sodium arsenite and 0.1 mM (2x) G3I compound (vortexed for 5 sec) was added to the sample. For heat shock experiments, each xy position was imaged every minute. At 2 min into the acquisition, the objective heater was ramped to 43°C. Following this, 250 μM (5x) G3I compound was prepared in FluoroBrite DMEM and vortexed for 15 sec. This solution was kept on a 47°C heater near the microscope along with a pipette tip. At 27 min into the acquisition (25 min into heat shock), 250 μL of the prewarmed solution was added to the cells using the prewarmed tip. We note that the solution was heated to 47°C to compensate for the rapid heat loss when adding the solution. At 32 min into the acquisition (30 min heat shock), the heat was ramped down to 37°C. All imaging was acquired on a Yokogawa CSU W1 spinning disk attached to a Nikon Ti2 eclipse with a Photometrics Prime 95B camera using Nikon Elements software (v5.20.00 to v5.21.02). Imaging was performed through a Nikon Plan Apo 60× 1.40 NA oil objective with Immersol 518 F (Zeiss; refractive index 1.518), and Perfect Focus 2.0 (Nikon) was engaged for all captures.

### Image analysis of pre-incubation and post-addition of compounds

Automated granule detection and measurement was performed using a combination of ilastik and CellProfiler software. Briefly, ND2 multipoint timelapse files were resaved as image sequences in Fiji with the Bio-Formats plugin so that individual frames of the movies could be treated as individual images for analysis. From there, pixel classification in ilastik was used to segment both stress granules and the total cellular area within the field using the G3BP1-GFP channel via machine learning. These segmentations, along with the original G3BP1-GFP image, were input into CellProfiler where objects were defined using the masks from ilastik. Using these data points, the measurement of the granules and total cellular area was performed.

### Immunofluorescence in cell lines

G3BP1-GFP cells were seeded in 4-well Lab-Tek II Chambered Coverglass (Thermo Scientific; 1055360). For pre-incubation experiments, cells were treated with 50 μM G3I compound for 20 min, followed by 250 μM NaAsO_2_ and 50 μM G3I compound for 30 min. For post-treatment, cells were treated with 250 μM NaAsO_2_ for 30 min, followed by treatment by 250 μM NaAsO_2_ and 50 μM G3I compound for 5 min. Cells were then washed twice with PBS and fixed with 4% PFA in PBS for 10 min. Cells were then washed 3 times with PBS (5 min between each wash), permeabilized with 0.5% Triton-X in PBS for 10 min, and washed twice with PBS. Next, cells were blocked for 1 h with 3% BSA in PBS at room temperature, followed by incubation with primary antibodies diluted in 3% BSA overnight at 4°C. Cells were then washed 3 times with PBS (5 min between each wash) and incubated with secondary antibodies and Hoechst diluted in 3% BSA at room temperature for 1 h. Cells were then washed 3 times with PBS (5 min between each wash) and imaged. Primary antibodies were as follows: eIF3η (Santa Cruz Biotechnology; sc-137214) at 1:100 dilution; PABP (Abcam; ab21060) at 1:1,000 dilution; and FXR1 (Abcam; ab51970) at 1:100 dilution. Secondary antibodies were as follows: donkey anti-mouse Alexa Fluor 647, donkey anti-rabbit Alexa Fluor 555, and donkey anti-goat Alexa Fluor 555 (Thermo Fisher Scientific). Nuclei were stained with Hoechst 33342 (Biotium; 40046) at 1:10,000 dilution. All imaging was acquired on a Yokogawa CSU W1 spinning disk attached to a Nikon Ti2 eclipse with a Photometrics Prime 95B camera using Nikon Elements software (versions 5.20.00 to 5.21.02). Imaging was performed through a Nikon Plan Apo 60× 1.40 NA oil objective with Immersol 518 F (Zeiss; refractive index 1.518), and Perfect Focus 2.0 (Nikon) was engaged for all captures.

### Imaging of mutation-induced granules and analysis

U2OS cells with knock-in of G3BP1-tdTomato [5] were seeded into 96-well plates (Corning; 3904). Either GFP-VCP-A232E or GFP-FUS-R495X were transfected into cells using ViaFect transfection reagent (Promega) 24 h prior to the start of the experiment. For samples to be treated with compounds G3Ia or G3Ia′, Hoechst (Biotium) was added and incubated for 30 min prior to the start of the experiment, washed once with PBS, and the media changed to 100 μL FluoroBrite DMEM media (Gibco) upon starting the experiment. Imaging of plates was performed on a Cytation C10 spinning disk confocal (BioTek) with a Hamamatsu Orca-Flash 4.0 camera using Gen5 software (version 3.11). Imaging was performed through an Olympus Plan Apo 40x 0.6 NA dry objective with adjustable collar set to 0.5 μm thickness. Laser autofocus prior to imaging the tdTomato channel was utilized for each image. The temperature was maintained at 37°C on the instrument and the cells were supplied with 5% CO_2_. A 7×7 tilescan was taken at the center of each well, capturing the tdTomato and GFP channels (and Hoechst when present). After completing the tilescan, 100 μL FluoroBrite with 100 μM (2x) G3I compound (vortexed for 5 sec) was added to each well and incubated for 20 min before starting another round of imaging in the same locations as the first round. Samples treated with G3Ib or G3Ib′ were analyzed manually, scoring each cell with spontaneous granules for the presence or absence of granules following treatment with compound.

Samples treated with G3Ia or G3Ia′ were analyzed using an automated pipeline. Segmentation was performed in both ilastik and Cellpose 2.0. Nuclear segmentation using the Hoechst channel and granule segmentation in both GFP and tdTomato channels was performed via pixel classification in ilastik using small cropped portions of the tilescan as the training dataset for each. Individual cell segmentation was performed in Cellpose 2.0 by inputting a merged RGB of the Hoechst and G3BP1-tdTomato channels. The Cellpose output was then eroded by 1 pixel and set as a binary mask in Fiji. The original 3 raw channels, 3 output masks from ilastik, and the 1 mask generated from Cellpose were all inputted into CellProfiler to generate per-cell measurements for granules as defined by GFP or tdTomato.

### Co-immunoprecipitation

U2OS cells expressing G3BP1-GFP were grown in a 3-cm dish to 100% confluency, at which point cells were collected and pelleted at 400 x g for 5 min. The pellet was either stored at −80°C or processed the same day by lysing with 200 μL in vivo lysis buffer (150 mM NaCl, 50 mM Tris [pH 7.5], 1 mM EDTA, 0.5% NP-40, 10% glycerol, supplemented with 1 protease inhibitor tablet (Roche 11836170001]). Lysates were centrifuged at 13,000 x g at 4°C for 15 min, after which supernatants were transferred to a new tube and diluted with 200 μL lysis buffer containing indicated concentrations of G3I compounds. The mixture was then incubated with GFP-Trap (ChromoTek GFP-Trap Magnetic Agarose; Proteintech; gtma) for 2 h at 4°C. Beads were washed three times with lysis buffer, then incubated with 1x SDS sample buffer supplemented with 10% 2-mercaptoethanol, and boiled for 5 min at 95°C. Lysates were subjected to SDS-PAGE and then immunoblotted.

### iPSC neuron differentiation

i3N iPSC clones were gifts from Michael E. Ward (NIH) and were used to generate a G3BP1-mNeongreen knock-in iPSC line. iPS neurons were differentiated with a two-step protocol (pre-differentiation and maturation) as previously described [41]. For pre-differentiation, iPSCs were incubated with 1 µg/ml doxycycline hyclate (Sigma Aldrich) for 3 days at a density of 1.2 x 10^6^ cells/well in six-well dishes coated with Matrigel in knockout DMEM (KO-DMEM)/F12 medium (Thermo Fisher Scientific) containing N2 supplement (Thermo Fisher Scientific), non-essential amino acids (NEAA; Thermo Fisher Scientific), GlutaMAX Supplement (Thermo Fisher Scientific), and Y-27632 (STEMCELL Technologies). The medium was [42]changed daily, and Y-27632 was removed from day 2. For maturation, pre-differentiated precursor cells were dissociated, counted, and sub-plated at 25 x 10^4^ cells/ml on dishes coated with 50 μg/ml poly-L-ornithine in BrainPhys neuronal medium (STEMCELL Technologies) containing N2 (Thermo Fisher Scientific), B-27, 20 ng/ml BDNF (PeproTech), 20 ng/ml GDNF (PeproTech), 500 µg/ml dibutyryl cyclic-AMP (Sigma Aldrich), 200 nM L-ascorbic acid (Sigma Aldrich), 1 µg/ml natural mouse laminin (Thermo Fisher Scientific), 1 μM Ara-C (Sigma Aldrich), and 1 µg/ml doxycycline hyclate. Half-medium was changed every other day.

### Immunocytochemistry in iPSC-derived neurons

DIV 21 human iPSC cortical neurons were co-incubated with compounds (50 μM or vehicle) and 500 μM NaAsO_2_ for 1 h, then immunostained as described previously [42] with modifications. In brief, after fixation with 4% PFA, neurons were blocked with PBS containing 5% normal goat serum and 0.1% Triton-X for 1 h. Primary antibodies were diluted with TBST containing 1% normal goat serum and 0.1% Triton-X. After overnight incubation of primary antibodies at 4°C, neurons were washed three times with TBST, two times with PBS, and incubated with Alexa Fluor 488, Alexa Fluor 555, and Alexa Fluor 633-conjugated secondary antibodies (Invitrogen) at a dilution of 1:1,000 for 2 h at room temperature. Images were captured using a Cytation microscope.

### Assembly and disassembly of stress granules using time-lapse live-cell microscopy in iPSC-derived neurons

DIV 21 human iPS cortical neurons were incubated with 500 μM NaAsO_2_ for 30 min, after which 50 μM compound or vehicle were added. Live cell imaging was performed using a Yokogawa CSU W1 spinning disk. A Yokogawa CSU W1 spinning disk attached to a Nikon Ti2 eclipse with a Photometrics Prime 95B camera using Nikon Elements software was used in time-lapse live-cell imaging. Imaging was taken using a 60× Plan Apo 1.4NA oil objective and Perfect Focus 2.0 (Nikon) engaged for the duration of the capture. During imaging, cells were maintained at 37°C and supplied with 5% CO_2_ using a Bold Line Cage Incubator (Okolab) and an objective heater (Bioptechs). To monitor the assembly and disassembly of stress granules, multipoint images over 5 xy fields for each condition per one replicate were taken with the 488-nm laser. Images were taken at each xy position every 1 min.

## Author contributions

R.L. and J.H. designed compounds. B.D.F., J.M., H.N., U.Y., J.W., J.H., J.D., W.H., K.W., M.W., C.L. designed and performed experiments. B.D.F., J.M., H.N., U.Y., J.W., H.J.K., J.D., W.H., K.W., M.W., C.L., and R.L. analyzed data. B.D.F, H.J.K., R.M., R.P., and J.P.T. drafted and revised the manuscript. R.M., and J.P.T. supervised the overall study.

## Acknowledgments

We thank Natalia Nedelsky for editorial assistance. We thank Eamon Comer for medicinal chemistry insights and discussion.This work was supported by the Howard Hughes Medical Institute and R35NS097974 grants to J.P.T.

